# Tissue Regeneration Requires Edema Fluid Clearance by Compensatory Lymphangiogenesis in Zebrafish

**DOI:** 10.1101/2024.05.30.596701

**Authors:** Olamide Olayinka, Hannah Ryu, Xiaowei Wang, Asrar B. Malik, Hyun Min Jung

## Abstract

Lymphangiogenesis is essential for tissue regeneration and edema clearance, and delayed or failed lymphangiogenesis is a critical cause of impaired healing. However, elucidating the dynamic changes in lymphangiogenesis during tissue regeneration in animal models in real-time has been challenging; thus, the mechanisms of compensatory lymphatic activation for edema clearance remain unclear. To address this, zebrafish were subjected to osmotic stress using a hypertonic solution (375 mOsm/L) and hypotonic solution (37.5 mOsm/L) to induce tissue damage followed by edema formation. Intravital imaging of *Tg(mrc1a:egfp; kdrl:mcherry)* larvae unveiled substantial lymphatic vessel remodeling during tissue regeneration. The increase of lymphatic endothelial progenitor cells accompanied by sustained expansion and remodeling of primary lymphatics during recovery suggested active lymphangiogenesis during tissue repair. We developed a novel method using translating ribosome affinity purification to scrutinize the translatome of lymphatic endothelial cells *in vivo*. This analysis revealed the upregulation of key pro-lymphangiogenic genes, notably vegfr2 and vegfr3, during tissue regeneration. Pharmacological inhibition of these Vegfr2/Vegfr3 prevented compensatory lymphangiogenesis and impaired healing. Our findings provide a new model of *in vivo* live imaging of regenerative lymphangiogenesis, presenting a novel approach to investigating lymphatic activation during tissue regeneration.

## INTRODUCTION

Tissue regeneration requires lymphangiogenesis, a process of lymphatic vessel formation essential for body fluid regulation and immune response, and delayed or failed lymphangiogenesis is a critical cause of impaired healing^1–4^. However, over several decades, the role of lymphangiogenesis in tissue regeneration has received little attention compared to the enormous interest in angiogenesis. Tissue injury causes extracellular matrix degradation and changes in transcapillary oncotic and hydrostatic pressure, favoring increased transcapillary filtration and increased endothelial permeability to protein due to inflammation^5–7^. If edema fluid is not properly cleared and swelling persists during tissue regeneration, lymphatic dysfunction can occur that may eventually cause systemic failure and permanent edema^8, 9^. Edema state depends on the ability to clear tissue fluid, and lymphatic vessels play a major role in this process^10^. The activation of the VEGF signal has been shown to generate promising revenue for regrowing lymphatic vessels^11–13^. Despite these advances in the role of lymphatics in regulating tissue fluid balance, questions such as the role of lymphatic endothelial progenitor cells and the nature and kinetics of the compensatory lymphatic mechanism during tissue regeneration have not been addressed.

The zebrafish *Danio rerio* provides an experimentally accessible and genetically tractable model for studying tissue regeneration and vascular biology^14, 15^. Zebrafish larvae are optically clear, facilitating high-resolution imaging, and various transgenic tools are available for visualizing and studying lymphatic vessel development and function^16–25^. As the zebrafish lymphatic system shares structural and functional similarities with the mammalian lymphatic network, the results can also be extrapolated to mammalian lymphangiogenesis^16–18, 20, 23–25^. Zebrafish and other freshwater teleosts have evolved mechanisms to regulate ion transport and maintain the hyperosmolarity of their interstitial fluid relative to their external environment^26^. Environmental stressors such as abrupt changes in salinity can disrupt their tightly controlled osmoregulation and cause tissue damage followed by fluid accumulation in extracellular spaces^27–29^. These advantages make the zebrafish a superb model system for studying the link between fluid homeostasis and lymphatics during tissue regeneration.

Development of tissue edema as in congestive heart failure in patients with ineffective fluid drainage due to the inability of lymphatics to remove tissue fluid^30^. This is also seen in post-surgical patients associated with defective lymphatic vessels^31^. However, mechanisms of how the lymphatic system regenerates during tissue repair and clears edema fluid remain incompletely understood. Here, we report the dynamic changes of lymphangiogenesis during both edemagenic and recovery phases of tissue regeneration that facilitate edema fluid clearance to demonstrate an integrated picture of the compensatory lymphatic expansion mechanism using live imaging in the zebrafish.

## RESULTS

### Live imaging of edemagenesis in zebrafish

Freshwater teleosts such as zebrafish have evolved mechanisms to regulate interstitial fluid homeostasis in response to environmental changes induced by water osmolarity^26^. Uncontrolled osmoregulation in response to highly saline conditions causes tissue damage and fluid accumulation in extracellular tissue space, the characteristic feature of edema^27–29^. To address the basis of edema formation and mechanisms of clearance of edema fluid, we established an optimized method of inducing quantifiable and reproducible edema (**Fig. 1A**). The embryos were initially placed in regular egg water (osmotic balance) until 2 dpf and then dechorionated before exposing them to a period of hypertonic stress (osmotic imbalance, 3x Danieau buffer, 375 mOsm/L) for 24 h. They were then returned to isotonic egg water (0.3x Danieau buffer, 37.5 mOsm/L), which triggered edema formation. Edema formation during 3-4 dpf was visualized using brightfield time-lapse imaging (**Supplementary Video 1**). We carried out eleven independent experiments with 20 larvae in each group and showed highly reproducible results of >90% successful edema formation in larvae from each experiment (**Fig. 1B**). Edema was visible at 4 dpf in the whole body as compared to animals not exposed to the osmotic stress (**Fig. 1C and Supplementary Fig. 1A**). The first site of observable edema was in the pericardial area followed by swelling of other parts of the body (**Fig. 1D**). Measurements of the 2-dimensional area on a lateral view of the larvae showed expansion of body parts, including the pericardial area (**Fig. 1D, 3.32-fold**), yolk area (**Fig. 1E, 1.60-fold**), and trunk area (**Fig. 1F, 1.13-fold**) at 4 dpf as compared to untreated siblings. The difference in head edema was not apparent at 4 dpf (**Fig. 1G**). The anatomic area measurement was normalized to fish length to account for inherent variations in developmental maturation and overall body size affected by osmotic stress^32^. The dorsal view of larvae showed prominent swelling surrounding the yolk area that was less obvious in the lateral views (**Fig. 1H**).

**Figure 1:**
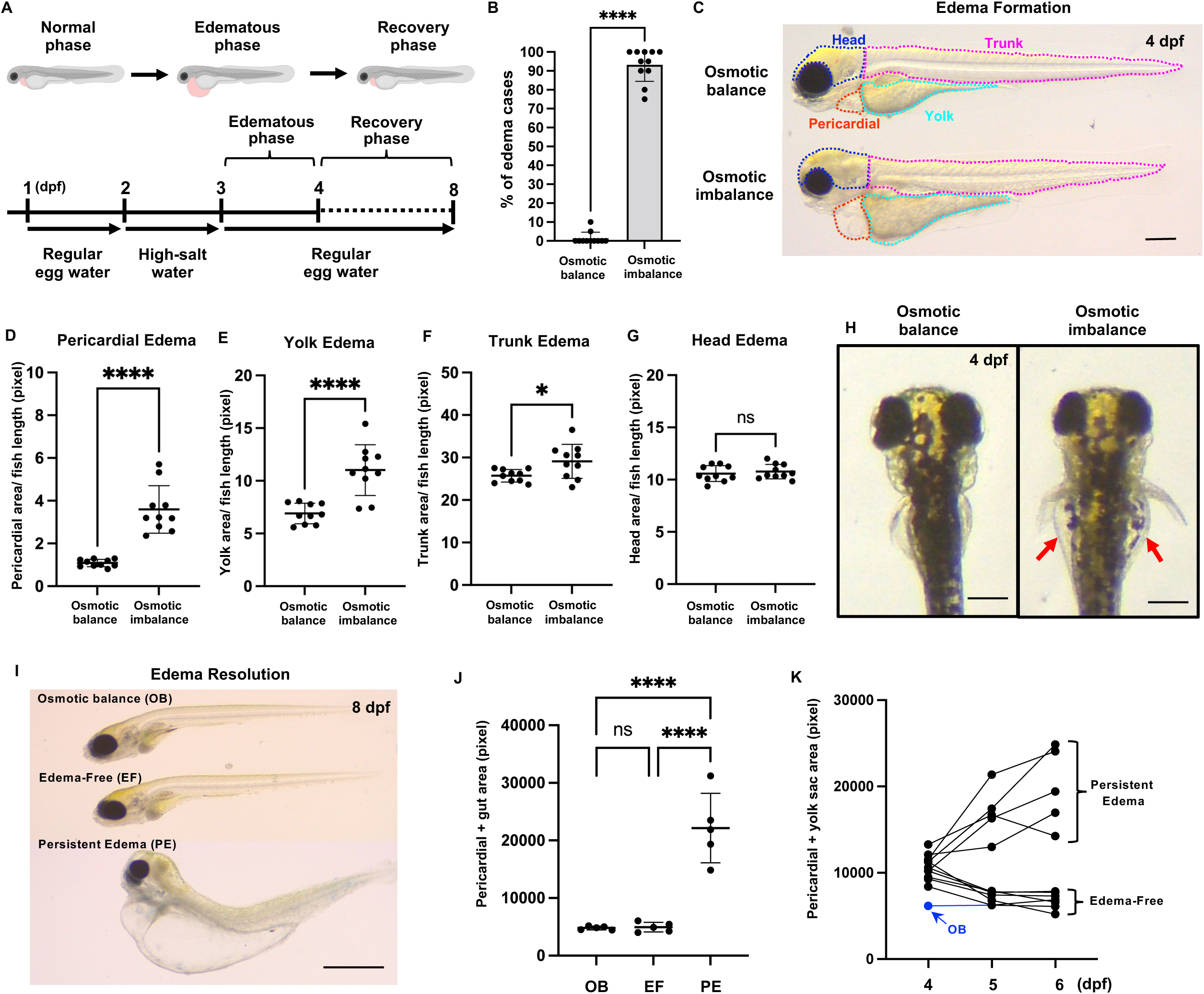
Zebrafish model of edema formation. (A) Schematic showing the experimental workflow from osmotic stress induction to edema formation and resolution. (B) Percentage of edematous larvae in osmotically balanced and osmotically imbalanced conditions at 4 dpf. Individual data points represent an independent experiment, and 20 larvae were used for each experiment. (C) Representative brightfield images showing lateral views of osmotically balanced (top) and osmotically imbalanced (bottom) 4 dpf zebrafish larvae. The dotted outline illustrates the boundary for edema area quantification in different parts of the body. (D-G) Quantification of pericardial area (D), yolk area (E), trunk area (F), and (G) head area. (H) Representative brightfield images showing the dorsal view of osmotically balanced (left) and osmotically imbalanced (right) zebrafish larvae. Red arrows indicate edema in the lateral side of the yolk area. (I) Representative images of osmotic balance (OB), edema-free (EF), and persistent-edema (PE) larva at 8 dpf. (J) Quantitative analysis of the pericardium and gut areas. Each dot represents an individual larva. (K) Line graph showing edema progression from 4-6 dpf. Each line represents the progressive change in the pericardial + yolk sac surface area of individual larvae over the experiment time course. The blue line is the larva raised in an osmotically balanced solution as a control. Scale bar = 500 μm (C,H,I). ns, not significant; *p<0.05; ****p<0.0001.

To study the time course of edemagenesis, we incubated larvae in the hypertonic solution for 18, 24, or 48 hours prior to osmotic stress. The number of edematous larvae increased when they were subjected to longer incubation in a hypertonic solution (**Supplementary Fig. 1B**). Notably, in a control experiment, prolonged incubation up to 48 h in the hypertonic solution in the absence of a return to isotonic water did not induce edema, indicating the need to switch from hypertonic state to isotonicity for edema formation (**Supplementary Fig. 1C,D**). Edema could also be induced when the protocol was delayed by 1 day (hypertonic exposure at 3-4 dpf followed by osmotic stress at 4-5 dpf), showing that this strategy can be used to target different developmental stages (**Supplementary Fig. 1E**). These results show that edema progress during tissue injury can be easily induced and imaged in real time in zebrafish larvae.

### Kinetics and reversibility of edema

To examine edema fluid clearance using this model, we next monitored the time course of changes in fluid accumulation. After a 24 h period of hypertonic stress followed by 24h of isotonic condition, we continued raising edematous larvae in regular isotonic egg water up to 8 dpf (**Fig. 1A**). We found that some larvae spontaneously resolved edema, whereas others developed persistent edema (**Fig. 1I**). The larvae with persistent edema displayed massive fluid accumulation in the whole body, including the head area where swelling was not apparent at earlier edema stage (**Fig. 1I**). In contrast, the edema-free larvae became morphologically indistinguishable from control larvae because the edema was completely resolved (**Fig. 1I,J**). Daily measurements of the edema progression in individual larvae showed that after the initial edema formed at 4 dpf, the first 24 hours (4-5 dpf) was the critical period determining the fate of larvae towards either recovery or remaining edematous (**Fig. 1K**). These data show that our new edema model provides an excellent system for visualizing edema formation and resolution during tissue regeneration.

### Generalized lymphatic vessel expansion as a characteristic feature of edema

Lymphatics are essential in recovery from edema and tissue healing by absorbing interstitial fluid from swollen areas^33, 34^. To address the changes of lymphatics in localized or global edema response, we next examined *Tg* (*mrc1a:egfp: kdrl:mcherry*) larvae and focused on the lymphatic vasculature changes in the trunk and craniofacial lymphatics at 8 dpf (**Fig. 2A**). At this developmental stage, control larvae are beginning to form lateral lymphatic vessels along the horizontal myosepta near where the transient parachordal lines (PL) had been present earlier in development (**Fig. 2B**). In comparison to the control osmotically balanced larvae (**Fig. 2B-D**), the edema-free “recovered” larvae displayed enhanced trunk lymphatic network formation, with more extended and enlarged TD, ISLVs, DLLV, and lateral lymphatics (LL) (**Fig. 2E-G**). We also observed particularly marked dilatation and remodeling of TD in the edema-free larvae (**Fig. 2G**). A 1.5-fold increase in the thoracic duct (TD) size in the edema-free group was seen as compared to the osmotically balanced control group (**Fig. 2H**). In addition, accelerated development of LL, with more somites containing LL segments at 8 dpf compared to controls following edema resolution (**Fig. 2I**).

**Figure 2:**
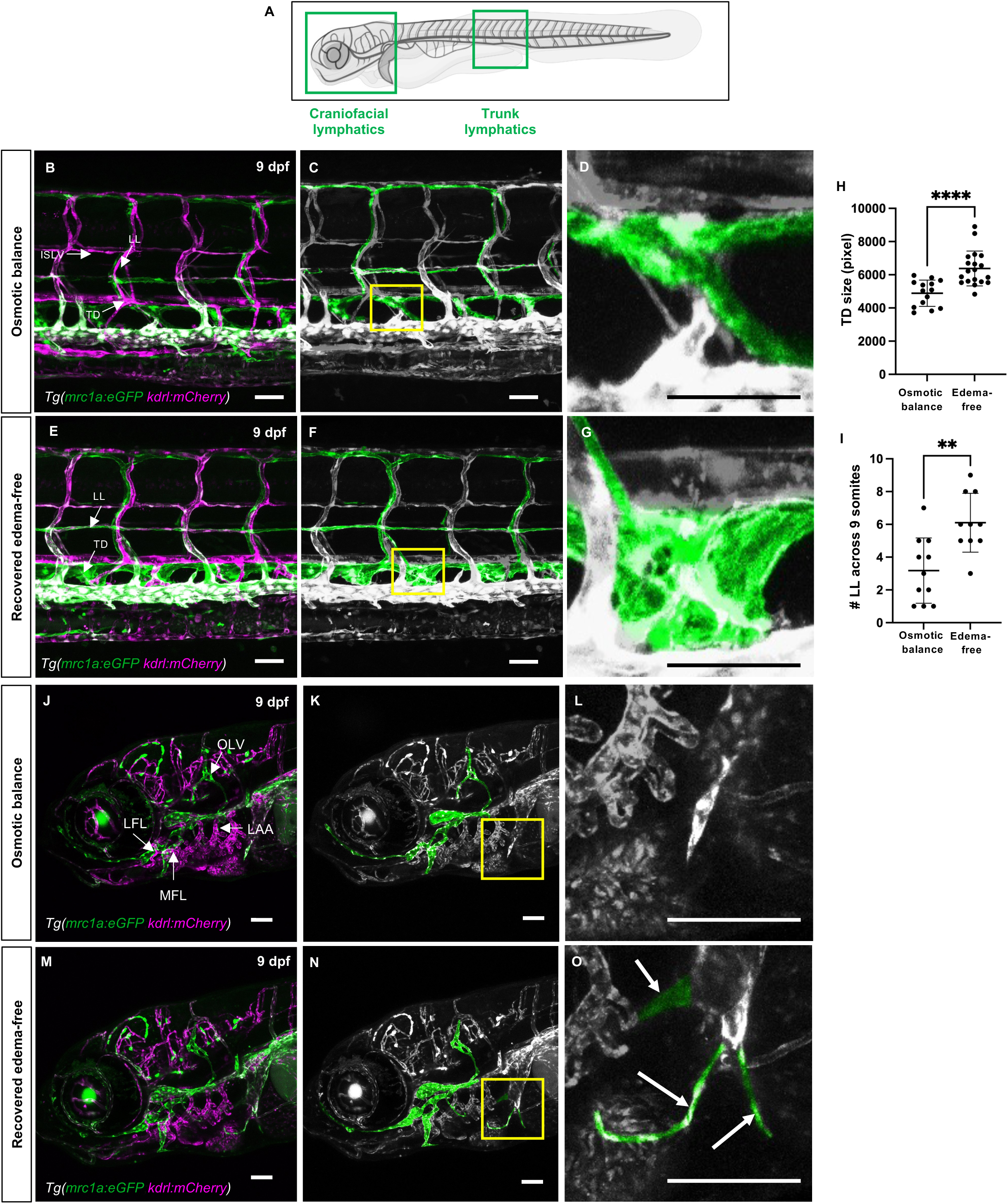
Lymphatic vessel expansion during edema recovery. (A) Schematic illustrating the imaging area to observe the lateral lymphatics and thoracic duct. (B) Confocal image of osmotically balanced *Tg(mrc1a:egfp;kdrl:mcherry)* larvae illustrating the normal vascular network at 9 dpf. (C) Lymphatic vessels are pseudo-colored in green from panel B. (D) Magnified image of the thoracic duct in panel C. (E) Confocal image of recovered edema-free *Tg(mrc1a:egfp;kdrl:mcherry)* larvae illustrating the vascular network following edema resolution. (F) Lymphatic vessels are pseudo-colored in green from panel F, highlighting the excess lateral lymphatics and dilated thoracic duct. (G) Magnified image of the thoracic duct in panel F. (H) Quantification of the thoracic duct surface area in osmotic balanced (n=14) and edema-resolved larvae (n=19). (I) Comparison of the number of LL segments in 9 somites between osmotic balanced (n=11) and edema-free larvae (n=9). (J) Confocal image of the facial lymphatics of osmotically balanced larvae at 9 dpf. (K) Lymphatic vessels are pseudo-colored in green from panel J. (L) Magnified image from panel K. (M) Confocal image of the facial lymphatics of recovered edema-free larvae at 9 dpf. (N) Lymphatic vessels are pseudo-colored in green from panel M. (O) Magnified image from panel N showing excessive lymphatic sprouts. Scale bar = 50 μm (B-G and J-O). **p<0.01; ****p<0.0001.

We also studied craniofacial lymphatics, which arise from lymphangioblasts in the common cardinal vein and primary head sinus at 1.5 dpf and anteroposteriorly migrate to form the lateral facial lymphatics (LFL)^20, 35^. Subsequent lymphangiogenesis forms medial facial lymphatics (MFL), otolithic lymphatic vessel (OLV), and lymphatic branchial arches (LAA) by 5 dpf from venous and non-venous origins^35^. We also examined the remodeling of craniofacial lymphatics at 9 dpf. As expected, control larvae maintained in the regular egg water had stereotypic facial lymphatic development (**Fig. 2J-L**). In contrast, edema-free larvae developed extended craniofacial lymphatics with larger LFL, MFL, and OLV (**Fig. 2M-O**). In addition, we found that edema-free larvae formed unusual lymphatic sprouts emerging from the common cardinal vein (**Fig. 2O and Supplementary Video 2**) and grew toward the direction of the pericardial space where there had previously been extensive edema (**Fig. 1**). These findings showed a broad activation of the lymphangiogenic program during tissue regeneration.

### Edema activates a lymphangiogenic program

To visualize lymphangiogenesis during osmotic stress-induced tissue repair, we imaged and quantitated the highly stereotypic lymphatic vasculature in the trunk of zebrafish larvae at the initiation of edema (**Fig. 3A**). The parachordal lines (PLs) are a transient population of lymphatic progenitors that localize to the horizontal myoseptum of the trunk at 2-3 dpf (**Fig. 3B**). The PL lymphangioblasts migrate dorsally and ventrally along trunk intersegmental arteries to form the intersegmental lymphatic vessels (ISLVs), thoracic duct (TD), and dorsal longitudinal lymphatic vessels (DLLVs), which together form the typical lymphatic network seen at 4-5 dpf (**Fig. 3B**)^16–18^. The control “osmotic balance” *Tg*(*mrc1a:egfp; kdrl:mcherry*) double transgenic animals showed typical initial lymphatic network patterns with ISLVs, TD, and DLLVs at 4 dpf (**Fig. 3C**), with only a few remnants of the PLs along the horizontal myoseptum at this stage (**Fig. 3D,E**). Although the edematous larvae also formed ISLVs, TD, and DLLVs (**Fig. 3F**), they had increased numbers of PLs at 4 dpf compared to controls (**Fig. 3G,H**). Quantitation of 9 somites anterior to the urogenital pore per larvae showed an overall 2-fold increase in PLs when edema is induced **(Fig. 3I)**. No changes in PL numbers were observed in the larvae incubated only with hypertonic solution for up to 48 hours, indicating the need for osmotic stress-induced edema to activate excessive production of PLs (**Supplementary Fig. 2A,B**).

**Figure 3:**
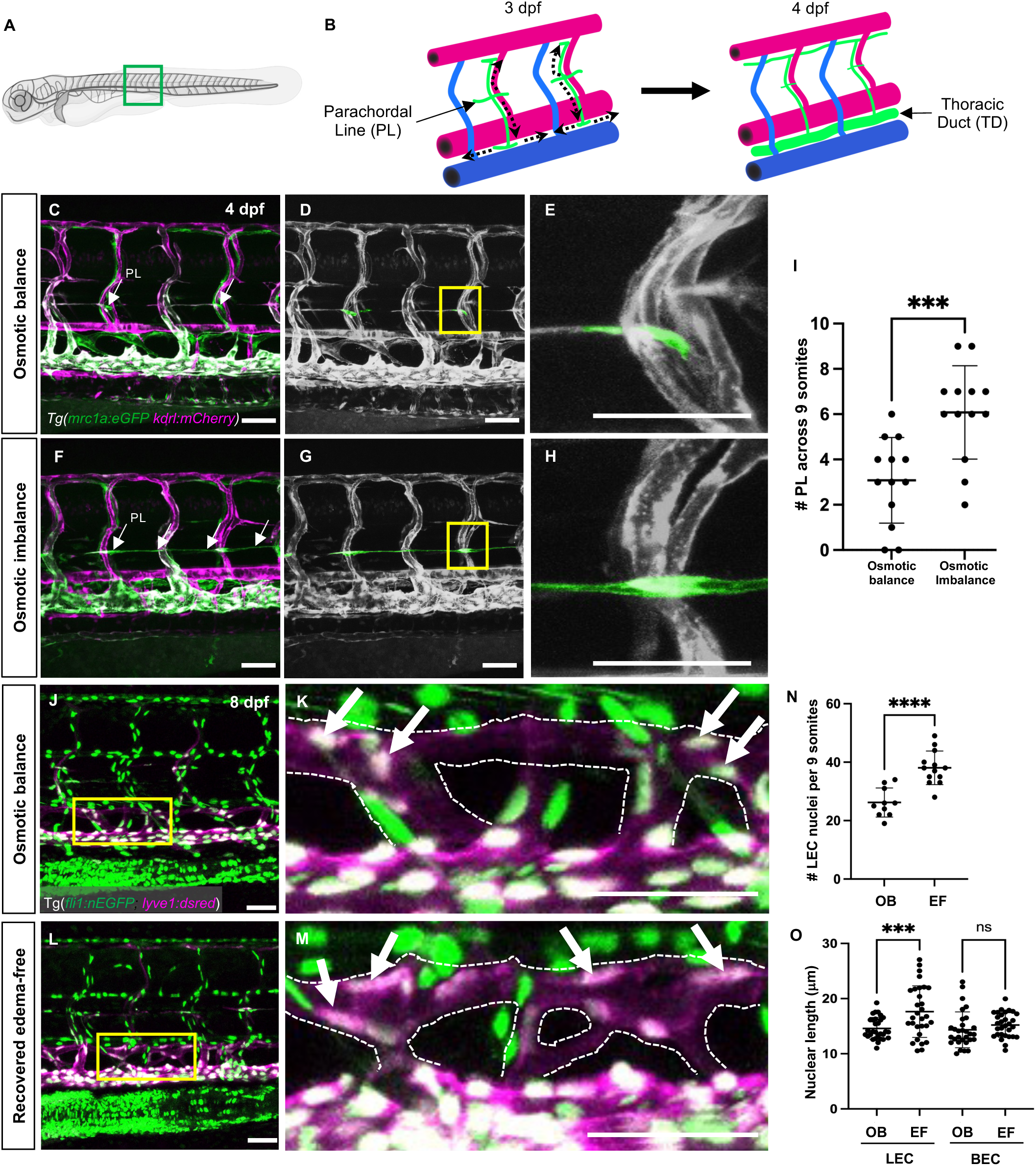
Lymphatic endothelial cell proliferation and elongation during edema recovery. (A) Schematic illustrating the imaging area of trunk lymphatics. (B) Diagram of developmental steps of lymphatic progenitors that give rise to lymphatic vessels at 3-4 dpf. (C) Confocal image of osmotically balanced *Tg(mrc1a:egfp;kdrl:mcherry)* larvae showing normal vascular network at 4 dpf. (D) Pseudo-colored confocal micrograph of untreated transgenic larvae indicating the parachordal line (lymphatic progenitors) in green and other vessels in grayscale. (E) Magnified image of the parachordal line in panel D. (F) Confocal image of osmotically imbalanced *Tg(mrc1a:egfp;kdrl:mcherry)* larvae showing the vascular phenotype in edema state. (G) Pseudo-colored confocal micrograph of osmotically imbalanced larvae highlighting the excess parachordal lines (lymphatic progenitors) during edema. (H) Magnified image of a parachordal line in panel F. (I) Quantification of the somites containing parachordal lines. 9 somites per larvae were counted (n=13 embryos each). (J) Confocal image of trunk vessels of an osmotically balanced *Tg(fli1:nEGFP;lyve1:dsred)* double transgenic larva. (K) Magnified image of the thoracic duct illustrating the oval-shaped LEC nuclei in panel J. The dotted lines indicate the boundary of the thoracic duct, and the arrows indicate LECs on the thoracic duct. (L) Confocal image of trunk vessels of a recovered edema-free *Tg(fli1:nEGFP;lyve1:dsred)* larva. (M) Magnified image of the thoracic duct highlighting the elongated LEC nuclei (arrow) in panel L. (N) Quantification of LECs in the thoracic duct of 9 somites between osmotic balanced (n=10) and edema-free larvae (n=13). (O) Quantification of lymphatic or blood endothelial nuclei length in osmotically balanced or edema-resolved larvae. A total of 30 nuclei were counted from each group. Scale bar = 50 μm (A-H and J-M). ns, not significant; ***p<0.001, ****p<0.0001.

To investigate changes in LEC number during edema and recovery, we analyzed the *Tg*(*fli1a;nEGPF), Tg(lyve1b:dsRed*) double-transgenic endothelial nuclear line with green endothelial nuclei and red lymphatic vasculature to visualize LEC nuclei and quantify the number of LECs in the lymphatic vasculature (**Fig. 3J-M**). Compared to the control osmotic-balanced larvae (**Fig. 3J,K**), the enlarged TD of edema-free recovered larvae had more LECs (**Fig. 3L,M**). Quantifying the number of TD LEC nuclei across 9 somites in 10 larvae from each group showed a 1.5-fold increase in LEC number in the edema-free recovered group compared to control osmotic-balanced larvae (**Fig. 3N).** We also observed significant elongation of LEC nuclei in edema-free larvae compared to controls (**Fig. 3J-M**). Interestingly, stretching of EC nuclei was observed only in LECs on the thoracic duct but not in the blood endothelial cells (BECs) in the adjacent posterior cardinal vein (**Fig. 3O**). These results together showed that the edema-induced pro-lymphangiogenic program was global as it involved massive remodeling and expansion of lymphatics.

### Endothelial translatome activation in edema-induced lymphangiogenesis

To probe the *in vivo* molecular changes in lymphatic endothelial cells in response to tissue damage and repair, we carried out transgene-assisted Translating Ribosome Affinity Purification (TRAP), which has been used to isolate ribosome-bound mRNAs undergoing active translation from specific cell types or tissues *in vivo*^36, 37^. This technique bypasses the cell dissociation steps that disrupt cell identity and alter gene expression, giving a true “*in vivo* snapshot” of molecular changes in highly specific cell populations^38–40^. To this end, we generated a novel *Tg(mrc1a:egfp-2a-rpl10a3xHA)* transgenic line using the mrc1a promoter^18^ to drive the expression of an eGFP-2A-rpl10a-3xHA transgene cassette^37^. This expression cassette consists of an eGFP coexpressed via a viral 2A peptide linkage with a 60S ribosomal protein l 10a (rpl10a) tagged with triplicate hemagglutinin epitope sequences (3xHA) to permit simultaneous visualization of transgene expression and translatome collection (**Supplementary Fig. 3A**). The “mrc1a-RiboTag” transgenic line is stably expressed in lymphatics and primitive veins, similar to a previously generated *Tg(mrc1a:egfp)^y251^* line (**Supplementary Fig. 3B**). We used this technique to profile the translatomes of lymphatic endothelial cells during edema progression. We collected control or osmotically stressed *Tg(mrc1a:egfp-2a-rpl10a3xHA)* larvae at 4 dpf, homogenized them to obtain crude lysates, and this was followed by affinity-purification of the HA-tagged ribosomes using anti-HA antibody and then direct immunoprecipitation to obtain the translating endothelial RNA (**Fig. 4A**). The control non-endothelial RNA was collected from the supernatant that contained untagged ribosome bound RNAs. We used TaqMan qRT-PCR to validate that the TRAP protocol enriched lymphatic and venous endothelial mRNAs. Gene expression analysis of flt4 (also known as vegfr3) and lyve1b, which are known to be expressed in lymphatics and veins at this developmental stage, were highly enriched in the endothelial RNA samples, indicating that the mrc1a-RiboTag transgenic line and the TRAP protocol work efficiently for analyzing endothelial translatome (**Supplementary Fig. 3C**). A ubiquitous gene n4bp1 (Nedd4 binding protein 1) showed no difference between endothelial and non-endothelial mRNA, reaffirming the specificity of this assay (**Supplementary Fig. 3C**). We used the *Tg(mrc1a:egfp-2a-rpl10a3xHA)* and TRAP-qPCR method to probe the changes in expression of key vascular genes potentially involved in responses to osmotic imbalance and edema. We induced edema (**Fig. 1A**), and then collected the edematous larvae and untreated siblings at 4 dpf. Subsequently, we used the TRAP protocol to obtain normal and edematous endothelial-specific mRNAs (**Fig. 4B**). We found that the expression levels of flt4, kdrl (also known as vegfr2), and lyve1b were significantly elevated in edematous larvae compared to the untreated siblings (**Fig. 4C**). The level of prox1a had an increasing trend but was not statistically significant, while vegfc levels were unchanged. Together, these data revealed the activation of flt4, kdrl, and lyve1b genes in lymphatic and venous endothelial cells as the signatures of the lymphangiogenic response during edema progression.

**Figure 4:**
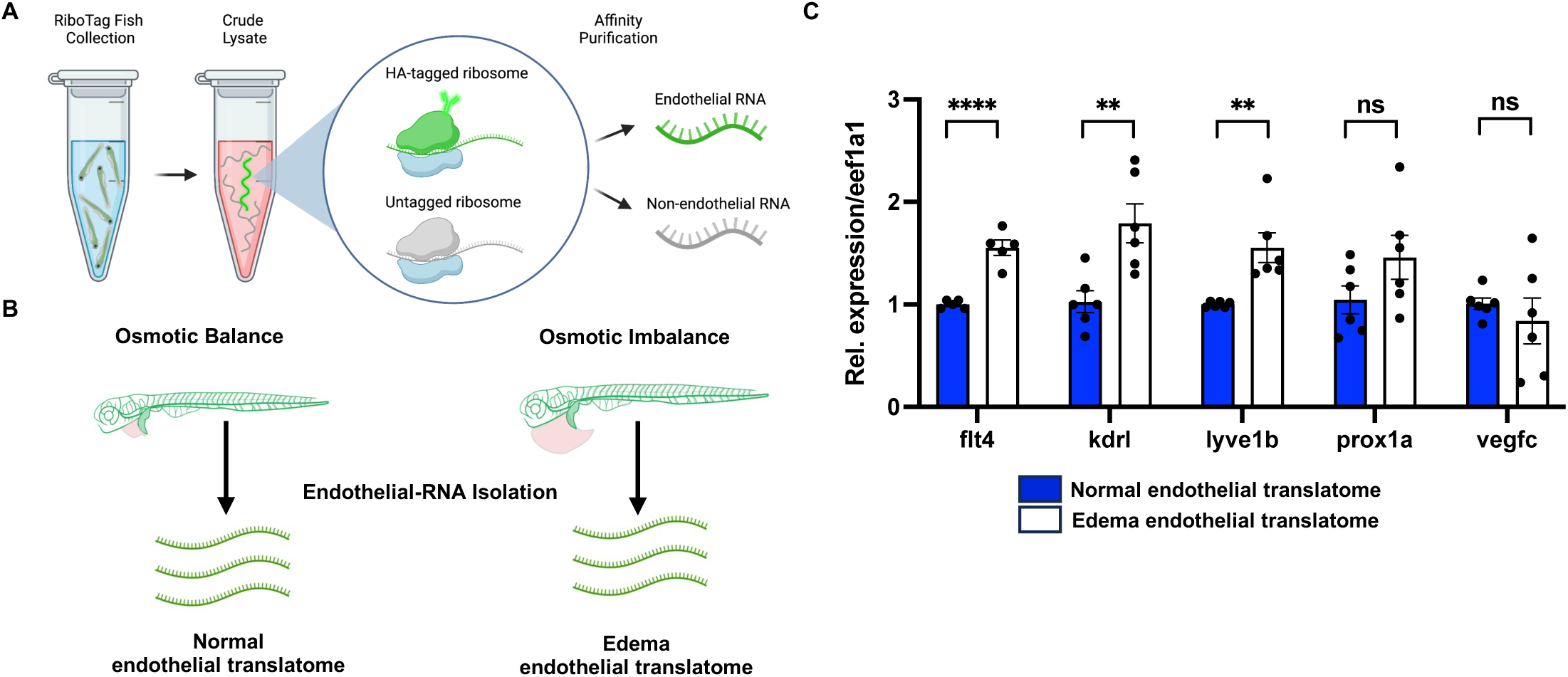
Endothelial translatome analysis reveals upregulation of lymphangiogenic factors in edematous vessels. (A) Schematic of the experimental flow of translating ribosome affinity purification (TRAP) protocol to isolate endothelial and non-endothelial translatome. (B) The experimental design to analyze endothelial translatome from osmotically balanced and imbalanced edematous Ribotag transgenic fish using TRAP assay. (C) Comparison of the translatome analysis of key lymphatic genes in osmotically balanced and imbalanced edematous larvae. Each RNA sample was obtained from a pool of at least 350 embryos. ns, not significant; **p<0.01; ****p<0.0001.

### Lymphatic insufficiency prevents edema fluid clearance

To examine whether the activation of lymphangiogenesis is required for edema recovery, we used the strategy of inhibiting the lymphangiogenic program. The Vegfr3 signaling pathway is essential for developmental and post-natal lymphangiogenesis^41, 42^. Similar to mammals, Vegfr3 (Flt4) signaling in the zebrafish mediates secondary sprouting from the posterior cardinal vein in the trunk and subsequent formation of horizontal myoseptum lymphatic progenitors (PLs) and ventral migration to form the thoracic duct^16, 17^. Since edema stimulates excessive PL formation (**Fig. 3**) and lymphatic expansion (**Fig. 2**), we examined whether this edema-induced lymphangiogenesis was Vegfr3-dependent using a Vegfr3 inhibitor SAR131675^43^. To specifically target excessive edema-induced PLs, we modified the edema induction protocol to begin the high-salt (3x Danieau) exposure at 72 hpf followed by edema induction with osmotic stress starting at 96 hpf (**Fig. 5A**). We treated larvae with Vegfr3 inhibitor SAR131675 starting at 78 hpf after secondary sprouts and PL formation was already underway (**Fig. 3B**). The larvae were maintained in regular egg water containing Vegfr3 inhibitor thereafter (**Fig. 5A**). The control larvae treated with DMSO displayed normal trunk and craniofacial lymphatic vessel phenotypes as expected (**Fig. 5B-E**). Larvae treated with Vegfr3 inhibitor at 78 hpf failed to form trunk lymphatics such as TD and ISLVs (**Fig. 5F,G**), but had only partially reduced MFL and OLV but no significant changes in the LFL and LAA (**Fig. 5H,I**). Although Vegfr3 is a central player in lymphangiogenesis, some lymphatic vessel beds also require Vegfr2, including the zebrafish craniofacial lymphatics^44^. We used SKLB1002 to inhibit Vegfr2^45^. Treatment of a combination of Vegfr2 and Vegfr3 inhibitors at 78 hpf prevented the development of TD and ISLVs in the trunk (**Fig. 5J,K**) as well as the craniofacial lymphatic branches that sprout after this developmental time point, including MFL, OLV, and LAAs (**Fig. 5L,M**). The FLF was found largely unaffected because it forms at an earlier stage prior to the inhibitor treatments^20, 35^. Modification of the protocol to inhibit Vegfr3 at a later time point was necessary because treatment of 10 μm Vegfr3 inhibitor SAR131675 to 22 hpf embryos caused complete loss of angiogenic sprouting and failed to form either intersegmental blood vessels or trunk lymphatic vessels in zebrafish larvae (**Supplementary Fig. 4A-E**). Similarly, the craniofacial lymphatic network development was also severely affected by defects in lateral facial lymphatics (LFL), medial facial lymphatics (MFL), lymphatic branchial arches (LAA), and otolithic lymphatic vessels (OLV) (**Supplementary Fig. 4F-I**).

**Figure 5:**
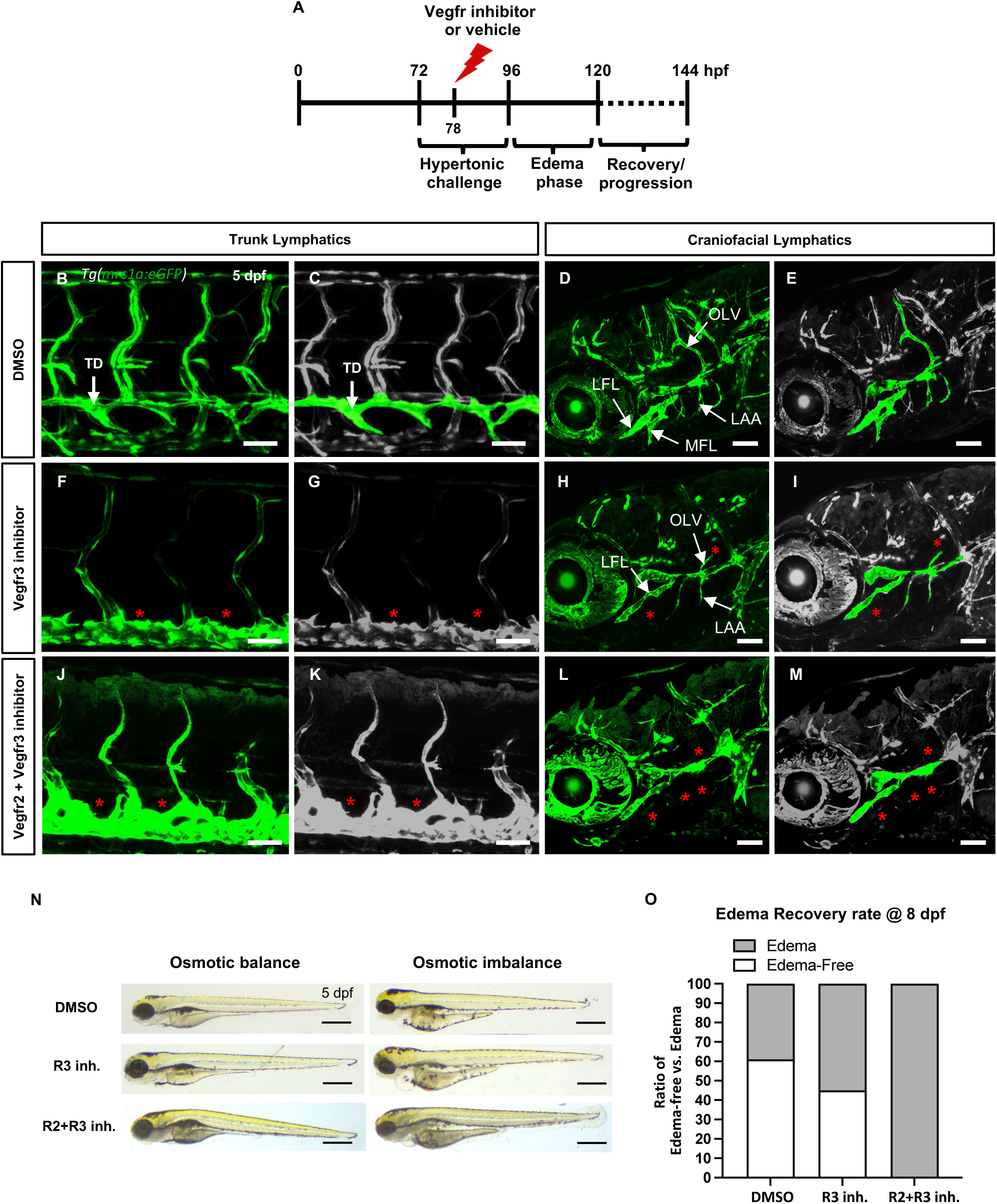
Vegfr-dependent lymphangiogenic program is required for edema fluid clearance. (A) Schematic diagram illustrating the experimental design for Vegfr inhibition during the osmotic challenge and edema progression. (B) Confocal image of trunk vessels of a DMSO-treated *Tg(mrc1a:egfp)* larva raised in an osmotically balanced condition. (C) Pseudo-colored confocal micrograph of a DMSO-treated larva highlighting the thoracic duct. (D) Confocal image of craniofacial vessels of a DMSO-treated *Tg(mrc1a:egfp)* larva raised in an osmotically balanced condition. (E) Pseudo-colored confocal micrograph of a DMSO-treated larva highlighting the facial lymphatics. (F) Confocal image of trunk vessels of a Vegfr3-inhibitor-treated *Tg(mrc1a:egfp)* larva showing complete absence of thoracic duct. (G) Pseudo-colored confocal micrograph of a Vegfr3-inhibitor-treated larva highlighting the loss of thoracic duct. (H) Confocal image of craniofacial vessels of a Vegfr3-inhibitor-treated *Tg(mrc1a:egfp)* larva showing partial absence of facial lymphatics. (I) Pseudo-colored confocal micrograph of a Vegfr3-inhibitor-treated larva highlighting the partial loss of facial lymphatics. (J) Confocal image of trunk vessels of a Vegfr2- and Vegfr3-inhibitor-treated *Tg(mrc1a:egfp)* larva showing complete absence of thoracic duct. (K) Pseudo-colored confocal micrograph of a Vegfr2- and Vegfr3-inhibitor-treated larva highlighting the loss of thoracic duct. (L) Confocal image of craniofacial vessels of a Vegfr2- and Vegfr3-inhibitor-treated *Tg(mrc1a:egfp)* larva showing absence of facial lymphatics. (M) Pseudo-colored confocal micrograph of a Vegfr2- and Vegfr3-inhibitor-treated larva highlighting the loss of facial lymphatics. (N) Representative brightfield images showing edema formation in 5 dpf larvae treated with DMSO, Vegfr3-inhibitor, or Vegfr2- and Vegfr3-inhibitors in osmotically balanced or osmotically imbalanced conditions. (O) The percentage of edema recovery demonstrated by the ratio of larvae recovered from edema or developed persistent edema at 8 dpf under DMSO and inhibitor treatments with osmotic stress. LFL, Lateral facial lymphatics; MFL, Medial facial lymphatics; LAA, Lymphatic brachial arches; OLV, Otholitic lymphatic vessels. Asterisks indicate missing lymphatic structures in panels F-M. Scale bar = 50 μm (B-M), 500 μm (N).

To address the requirement of lymphatics in edema fluid clearance, we induced edema using osmotic stress and subsequently prevented Vegfr pathways using inhibitors. Interestingly, despite the efficient blockage of trunk and craniofacial lymphatics with Vegfr2 and Vegfr3 inhibitors for 42h (78 hpf to 120 hpf), the edema was not produced in an osmotically balanced condition, indicating that lymphatics are dispensable in steady-state condition (**Fig. 5N**, **left panels**). Larvae exposed to osmotic stress produced edema regardless of the presence of lymphatics as treatment with DMSO, Vegfr3 inhibitor alone, or Vegfr2 and Vegfr3 inhibitors showed similar edema phenotype (**Fig. 5N**, **right panels**). We observed a dramatic difference in the edema recovery rate among these groups. About 60% of DMSO-treated edematous larvae under osmotic stress at 5 dpf recovered and became edema-free at 8 dpf (**Fig. 5O**). This recovery rate dropped by 15% in the Vegfr3 inhibitor-treated group, which showed partially impaired facial lymphatics and complete loss of trunk lymphatics (**Fig. 5F-I**). Strikingly, this rate was further reduced to 0% when larvae almost completely lost both trunk and craniofacial lymphatics by a combination of Vegfr2 and Vegfr3 inhibitors (**Fig. 5O**). All larvae with lymphatic insufficiency under osmotic imbalanced condition progressed to a persistent edema state and eventually died. These results showed that lymphatics play a central role in fluid clearance to reduce edema burden in zebrafish larvae.

## DISCUSSION

In this study, we address fundamental questions regarding the role of compensatory lymphangiogenesis in draining excess fluid accumulating during tissue regeneration. We present a new model for regenerative lymphangiogenesis during osmotic stress-induced tissue damage and investigate the essential role of lymphatic vessels in edema fluid clearance. Using this model, we demonstrate that regenerative lymphangiogenesis requires the activation of a lymphangiogenic program to produce lymphatic endothelial progenitors, followed by compensatory lymphatic expansion during the edema recovery process. We use a novel transgenic tool for endothelial translatome and show that Vegfr2 and Vegfr3 levels elevate by edema induction. Furthermore, we show that both VEGF receptors are required for regenerative lymphangiogenesis during the recovery. Overall, the edema recovery assay confirms the requirement of both Vegfr2 and Vegfr3 that are required for craniofacial lymphangiogenesis in the zebrafish.

The edema-induction model described in Methods allows quantifiable and reproducible edema formation. Since a single pair of zebrafish breeders produce 200-300 eggs, this approach also provides a statistically sound assessment as an experiment material^46^. The protocol is designed to align edema production with the time of natural lymphatic development. The lymphatic progenitors emerge from the posterior cardinal vein around 2 dpf and start to migrate to their designated locations^16–18^. The embryos were challenged with osmotic stress when the lymphatic progenitors actively propagate and migrate to their designated locations. The full recovery of edema was determined within 48 hours after the larvae were exposed to osmotic stress, which was accompanied by the expansion of the lymphatic network. Hypertonic stress induces cellular shrinkage and disruption of the epithelial junctional barrier, allowing water to be flooded into the interstitial space^47, 48^. Studies showed that long-term (∼2 weeks) supplement of a high-salt diet in rodents induces lymphangiogenesis via macrophage-secreted Vegfc and VEGFR3 signaling^49–51^. The relatively short duration (∼2 days) of hypertonic stress exposure to zebrafish larvae does not alter lymphangiogenesis. This indicates that the fluid accumulation created by the osmotic imbalance is the key driver in activating lymphatic vessel expansion rather than the salt treatment itself in this experimental setting. This is consistent with the observation seen in interstitial fluid accumulation in *ex vivo* mice embryo stimulating lymphatic endothelial cells (LEC) activation and lymphatic vessel expansion^52^, suggesting an evolutionarily conserved mechanism for pumping fluid from tissue. The lymphatic expansion was ascribed to the expression of Vegfc and matrix metalloproteinases^53^; however, the causal relationship between the signals and generation of *de novo* LEC and lymphatic vessels capable of clearing edema was not established.

Additional biological insights from this study come from our analysis of high-resolution intravital live imaging of regenerative lymphangiogenesis in zebrafish. This model allows analysis of a robust generation of lymphatic progenitor cells followed by extensive remodeling of lymphatic vessels during tissue regeneration. Several animal models have been developed to investigate the role of lymphatic vessels in mediating interstitial fluid drainage and edema clearance. Although the commonly used surgical model of lymphatic injury provides valuable information to tackle questions of lymphatic flow regulation^54, 55^, it is technically challenging to visualize lymphatic vessel regeneration because it greatly limits the spatial and temporal windows available to investigate edema progression and lymphatic changes, precluding a more holistic analysis, as reported by several studies^55–57^. The use of the zebrafish model overcomes this problem. It provides access to investigate the entire process of lymphangiogenesis during tissue regeneration, as shown by the initial changes during the edema formation phase until the full recovery phase.

Unexpectedly, ablating both craniofacial and trunk lymphatics in larvae did not produce edema under steady-state conditions when fluid balance was not perturbed. This indicates that other fluid regulatory organs, such as skin epithelium and pronephros, are sufficient for fluid homeostasis during steady-state conditions in early-stage larvae. However, lymphatics become essential when edema is present. Complete ablation of craniofacial and trunk lymphatics together drastically reduced the edema recovery rate to 0%. When the lymphatic is absent, no animals are able to recover from induced edema and die with persistent edema present. Our data also suggests that craniofacial lymphatics play a major role in removing excessive fluid at these developmental stages^58^. The tissue fluid-driven signal that activates the lymphangiogenic program is unknown. Accumulation of fluid in the interstitial space increases the interstitial hydrostatic pressure^59^; thus, it is possible that the activation of tissue pressure sensors such as Piezo1/2 or members of the *Trpc* family may be responsible.

Together, we report a valuable zebrafish model that provides an excellent *in vivo* system for live imaging edema progression and investigating the compensatory lymphatic expansion mechanism during tissue regeneration. We believe the work presented here provides important tools and information to launch future studies to dissect the cooperation between lymphatics, fluid homeostasis, and tissue regeneration.

## METHODS

### Zebrafish maintenance and fish strains

Zebrafish husbandry and research protocols were reviewed and approved by the University of Illinois Animal Care and Institutional Biosafety Committee, Animal Care and Use Protocol 23-112. All methods were performed in accordance with the relevant guidelines and regulations and conform to the ARRIVE guidelines^60^. Zebrafish embryos were obtained by natural spawning and raised in blue water (60 mg of Instant Ocean salt mix, 1 mL of 0.1% methylene blue per liter of RO water) at 28.5°C following standard husbandry conditions. *Tg(mrc1a:eGFP)^y251^*; Tg(kdrl:mCherry)^*y171*^ double transgenic line were used for the imaging studies^18^. *Tg*(*fli1:nEGFP)^y7^*; Tg (*lyve1:dsred)* double transgenic line was used in the quantification of lymphatic endothelial cell (LEC) numbers ^16, 20^. *Tg(mrc1a:egfp-2A-rpl10a-3xHA)* was used to isolate the endothelial translatome for quantitative PCR.

### Induction of edema with osmotic stress

Embryos were dechorionated at 48 hpf either manually or using 1 mg/mL pronase (Sigma-Aldrich) in blue water for 2-5 min. Dechorionated embryos were washed three times in fresh blue water to remove the remaining pronase in the solution. The embryos were incubated in hypertonic salt solution (3x Danieau buffer - 58 mM NaCl, 0.7 mM KCl, 0.4 mM MgSO_4_, 0.6 mM Ca(NO_3_)_2_, 10 mM HEPES) with an osmolarity of 375 mOsm/L for a period of 24 h (until 72 hpf) before being abruptly returned to isotonic solution (37.5 mOsm/L, 0.3x Danieau). This period of osmotic stress produced reproducible edema, which was quantified using the ImageJ free hand selection option to trace the boundary of the pericardial and yolk sac pouch from the lateral view at indicated developmental stages. The edema surface area was measured in pixels. Thoracic duct size was quantified by free hand tracing of the thoracic duct across 9 somitic segments rostral to the urogenital pore. Parachordal lines and lateral lymphatics were quantified by counting the number of somites with lymphatics in 9 somitic segments rostral to the urogenital pore. Surface area measurements were presented in area pixels.

### Studying effects of Vegfr inhibition

To test the effects of Vegfr inhibitors on the development of blood or lymphatic vessels in the presence or absence of edema, embryos were subjected to the osmotic challenge starting at 3 dpf and then treated with either Vegfr3 inhibitor SAR131675 (Selleck Chemicals) and/or Vegfr2 inhibitor SKLB1002 (Selleck Chemicals) or DMSO (as control) at indicated time points and incubated up to 5 dpf. The inhibitors were washed out at 5 dpf, and vascular changes during edema and resolution were quantified.

### Generation of RiboTag construct and transgenic lines

The RiboTag construct was generated using Tol2kit components with Gateway Technology^61^. To generate mrc1a:eGFP-2A-rpl10a-3xHA DNA construct, LR clonase II (Thermo Fisher Scientific) was used to recombine p5E-mrc1a promoter^18^, pME-eGFP-2A-rp110a-3xHA^37^, and p3E-polyA^61^ into pDestTol2pA2^61^. The DNA construct and Tol2 transposase RNA were microinjected into the blastomere of one-cell stage zebrafish embryos. A stable *Tg(mrc1a:egfp-2a-rpl10a-3xHA)* germline was established by screening through multiple generations. This transgenic line isolates the active lymphatic and venous endothelial translatome at desired developmental stages.

### Translating Ribosome Affinity Purification (TRAP)

TRAP was performed following a modified protocol^37^. For each untreated or edematous sample, 500 4dpf mrc1a RiboTag zebrafish were dechorionated, deyolked, and dounce-homogenized in homogenization buffer consisting of 1 mM Tris pH 7.4, 100 mM KCl, 12 mM MgCl_2_, 1% NP-40, 1 mM DTT, 1x Protease inhibitors (Sigma), 200 units/mL RNAsin (Promega), 100ug/mL cycloheximide (Sigma) and 1mg/mL heparin (Sigma). Lysates were incubated on ice to ensure complete cell lysis and then cleared for 10min at 10,000xg, 4^0^C. 4 µl of anti-HA antibody (Abcam ab9110 Rabbit polyclonal) was added per 500 embryos in 800µl lysate, and the samples were orbitally rotated at 4^0^C for 4h. 100 µl of Dynabeads® protein G slurry (Novex/Life Technologies) was added per 1 µl of anti-HA antibody with homogenization. Dynabeads were pre-washed with 800 ul homogenization buffer, rotating for 30min to equilibrate the beads. The wash buffer was then removed, and the lysate + Ab solution was added to the equilibrated Dynabeads. This dynabead + lysate + Ab mixture was incubated overnight at 4^0^C in an orbital rotator. Dynabeads were collected using a magnetic stand and washed three times for 5m each in an orbital shaker with high salt homogenization buffer (50 mM Tris pH 7.4, 300 mM KCl, 12 mM MgCl_2_, 1% NP-40, 1mM DTT, 1x Protease inhibitors, 200 units/mL RNAsin, 100ug/mL cyclohexamide, 1mg/mL heparin). Polysome-bound RNA was retrieved from Dynabeads and DNase-treated using the Direct-zol^TM^ RNA MicroPrep kit (Zymo Research).

### Image acquisition and processing

Embryos were anesthetized using 1x tricaine (MS-222) in RO water and laterally or dorsally mounted in 1% low melting point agarose gel, placed on a 35 mm dish with No. 1.5 coverslip and 14 mm glass diameter. Stereoscope images were obtained using the Zeiss Stemi 305 microscope equipped with Axiocam Erc 5s camera (Zeiss) and EVOS M5000. Edema formation timelapse video was obtained in mp4 format at 5 frame rates per second (fps) using EVOS M5000. Confocal microscopic imaging was performed with the Zeiss LSM 880 confocal microscope. High-resolution z-stack images were obtained with 10x or 20x objective lenses at 1024 x 1024 frame size. The bi-directional image scanning speed was set at 7. Image analysis was done using ImageJ (version: 2.3.0/1.53q; Build: d544a3f481) and Adobe Photoshop (Adobe).

### Statistical analysis

Statistical analysis was determined using an unpaired two-tailed student’s t-test for pair-wise comparison and a one-way ANOVA test for group comparison. Data are presented as mean ± SD. Significance was established when the p-value was under 0.05. Analysis was performed using GraphPad Prism Software, version 9.2.0.

## ACKNOWLEDGMENTS

We are grateful to UIC Biological Resource Laboratory for their assistance in zebrafish facility care. We also thank Ms. Michelle Dharma at UIC for her assistance in some of the experiments and her excellent care of the zebrafish used in this study. This research was supported by the American Heart Association (grant number 24PRE1199881) to O.O. and the UIC College of Medicine start-up fund to H.M.J.

## AUTHOR CONTRIBUTIONS

OO, HR, and HMJ performed the experiments and analyzed data. OO, ABM, and HMJ contributed to the conceptualization and experimental design. OO, XW, ABM, and HMJ contributed to writing, reviewing, and editing the manuscript. HMJ conceived the study and supervised the work. All authors have read and agreed to the published version of the manuscript.

## DATA AVAILABILITY

All data that support the findings of this research are included in the article and the supplementary files.

---

**Supplementary Figure 1.**
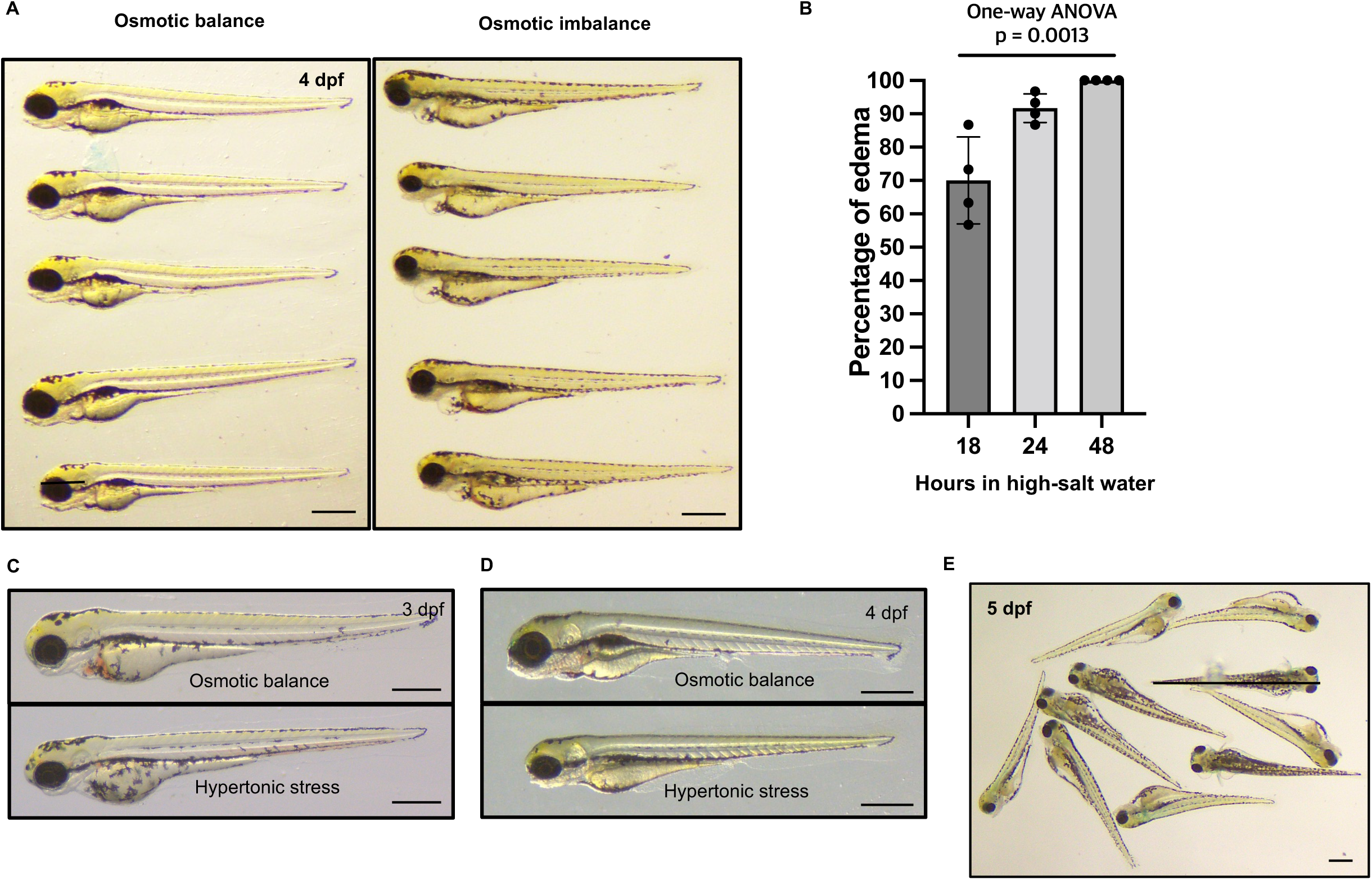
Zebrafish edema model. (A) Representative brightfield images of five osmotically balanced and imbalanced 4 dpf zebrafish larvae in a lateral view. (B) Percentage of edematous larvae produced by the edema induction protocol using 18, 24, or 48 hours of incubation in hypertonic solution. Four dots represent independent experiments with 30 larvae. (C,D) Brightfield images showing osmotic balanced larvae and larvae subjected to hypertonic stress at 3 dpf (B) and 4 dpf (C). (E) Brightfield group image of larvae subjected to the modified edema induction protocol with incubation in hypertonic solution for 48 h. Scale bars = 500 μm (A, C-E).

**Supplementary Figure 2:**
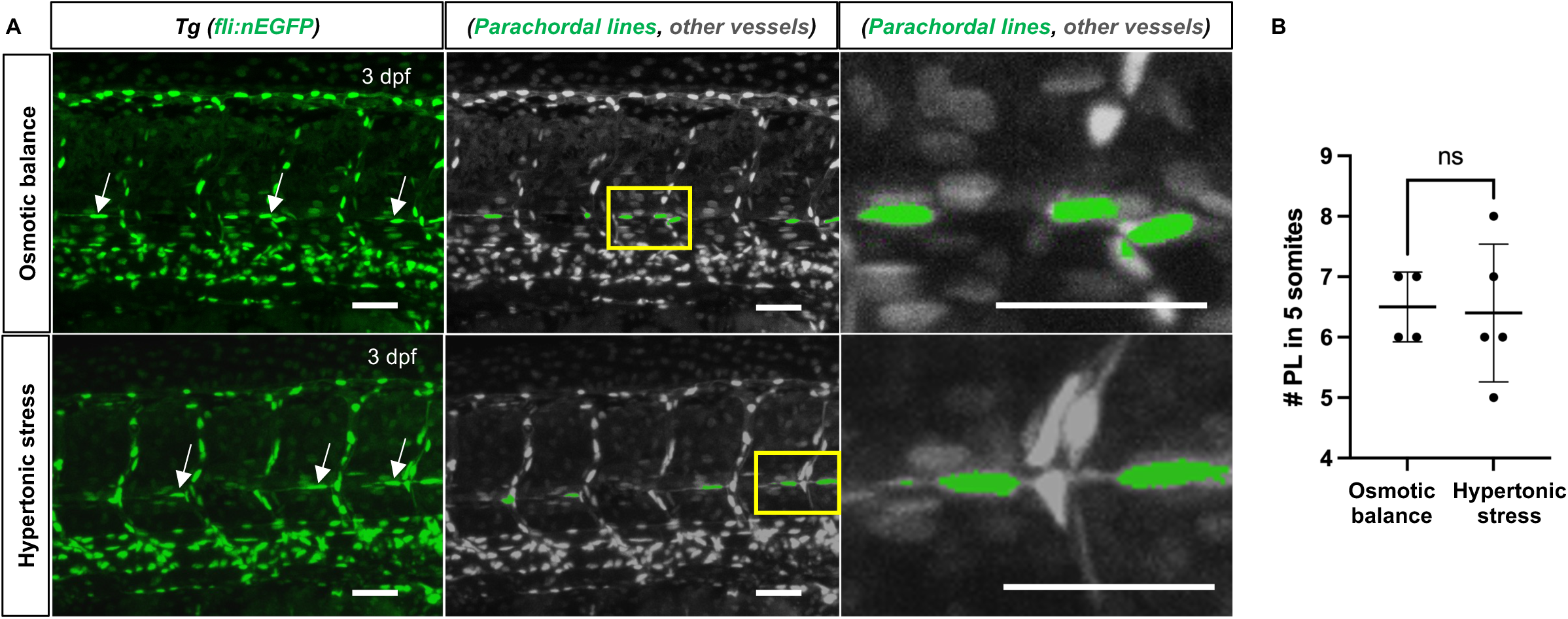
Hypertonic stress alone does not induce lymphatic expansion. (A) Confocal images of 3 dpf *Tg(fli1:nEGFP)* transgenic larvae raised in an osmotically balanced or hypertonic solution condition for 24 hours. (B) Quantification of parachordal line formation in larvae raised in osmotically balanced or hypertonic solution for 24 hours. Scale bars = 50 μm (A). ns, not significant.

**Supplementary Figure 3:**
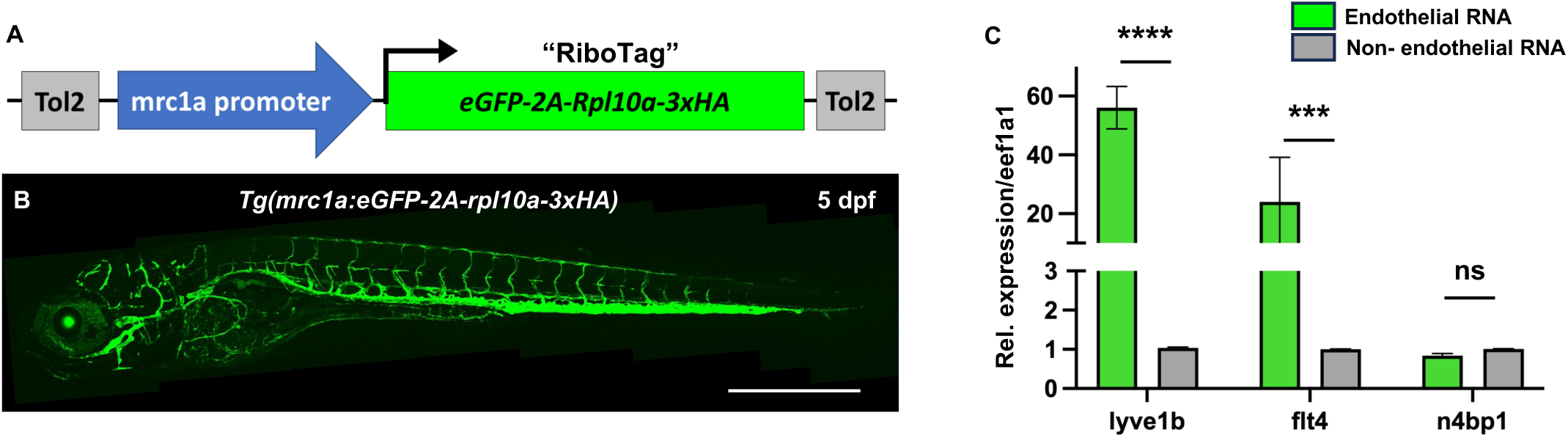
Lymphatic and venous translatome enrichment using mrc1a:RiboTag transgenic animal. (A) Schematic diagram illustrating the mrc1a Ribotag DNA construct. (B) Representative image of the stable transgenic line *Tg(mrc1a:eGFP-2A-rpl10a-3xHA)* at 5 dpf. (C) qPCR data showing the relative expression of lyve1b and flt4 enriched in endothelial translatome compared to the non-endothelial pool. The n4bp1 expression was used as an example of a non-endothelial gene. Each RNA sample was obtained from a pool of at least 350 embryos. Scale bar = 500 μm (B). ns, not significant; ***p<0.001, ****p<0.0001.

**Supplementary Figure 4:**
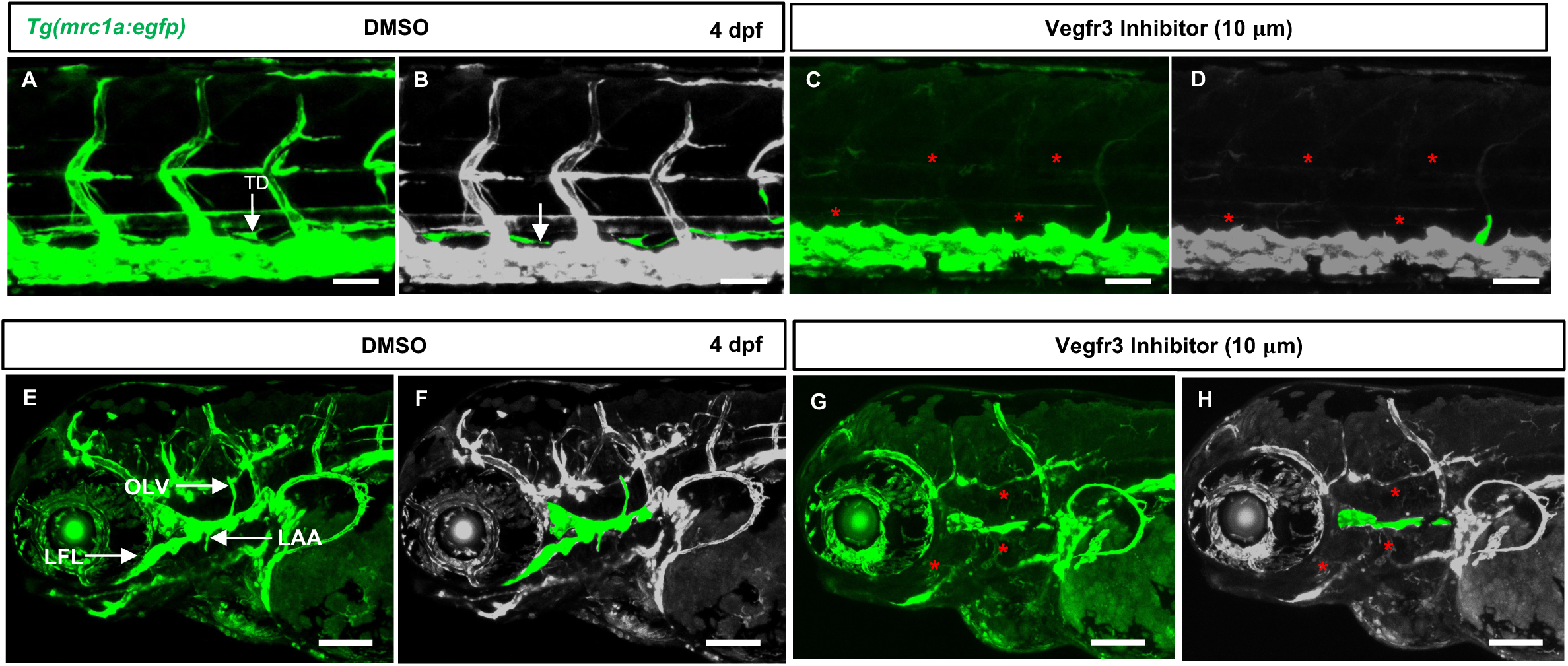
Early vegfr3 inhibition significantly ablates lympho-venous sprouting and disrupts both trunk and craniofacial lymphangiogenesis. (A) Confocal image of trunk vessels in a DMSO-treated *Tg(mrc1a:egfp)* larva at 4 dpf. (B) Pseudo-colored image of panel B highlights the thoracic duct in green. (C) Confocal image of a Vegfr3-inhibitor-treated *Tg(mrc1a:egfp)* larva showing complete ablation of trunk lymphatics. (D) Lymphatics are pseudo-colored in green. (E) Confocal image of craniofacial vessels of a DMSO-treated *Tg(mrc1a:egfp)* larva at 4 dpf (F) Pseudo-colored image of panel F highlights the early facial lymphatic structures in green. (G) Confocal image of craniofacial vessels of a Vegfr3-inhibitor-treated *Tg(mrc1a:egfp)* larva showing defective facial lymphatics. (H) Pseudo-colored image of panel H highlights the absence of facial lymphatic structures. Scale bar = 50 μm (A-H).

**Supplementary Video 1: Live imaging of edema formation.**

A timelapse video of an osmotically balanced larva (top) and an osmotically imbalanced larva (bottom) illustrating edema formation. A total of 160 frames of time-lapse images were collected from 96 hpf to 120 hpf every 10 minutes. The time-lapse movie was created at 65 frames per second. The arrows indicate the fluid accumulation in the osmotically imbalanced larva. Scale bar = 200 μm.

**Supplementary Video 2. 3-D movie showing craniofacial vessels in edema-free larva.**

3-D image reconstruction of the craniofacial vessels of an edema-free 9 dpf *Tg(mrc1a:eGFP; kdrl:mCherry)* double-transgenic larva, highlighting lymphatic sprouts directed toward the pericardial area.

